# Arabidopsis JASMONATE-INDUCED OXYGENASES down-regulate plant immunity by hydroxylation and inactivation of the hormone jasmonic acid

**DOI:** 10.1101/102103

**Authors:** Lotte Caarls, Joyce Elberse, Mo Awwanah, Nora R. Ludwig, Michel de Vries, Tieme Zeilmaker, Saskia C.M. Van Wees, Robert C. Schuurink, Guido Van den Ackerveken

## Abstract

The phytohormone jasmonic acid (JA) is vital in plant defense and development. Although biosynthesis of JA and activation of JA-responsive gene expression by the bioactive form JA-isoleucine (JA-Ile) have been well-studied, knowledge on JA metabolism is incomplete. In particular, the enzyme that hydroxylates JA to 12-OH-JA, an inactive form of JA that accumulates after wounding and pathogen attack, is unknown. Here, we report the identification of four paralogous 2-oxoglutarate/Fe(II)-dependent oxygenases in *Arabidopsis thaliana* as JA hydroxylases and show that they down-regulate JA-dependent responses. As they are induced by JA we named them *JASMONATE-INDUCED OXYGENASEs* (*JOXs*). Concurrent mutation of the four genes in a quadruple Arabidopsis mutant resulted in increased defense gene expression and increased resistance to the necrotrophic fungus *Botrytis cinerea* and the caterpillar *Mamestra brassicae*. In addition, root and shoot growth of the plants was inhibited. Metabolite analysis of leaves showed that loss of function of the four JOX enzymes resulted in over-accumulation of JA and in reduced turnover of JA into 12-OH-JA. Transformation of the quadruple mutant with each *JOX* gene strongly reduced JA levels, demonstrating that all four JOXs inactivate JA in plants. The *in vitro* catalysis of 12-OH-JA from JA by recombinant enzyme could be confirmed for three JOXs. The identification of the enzymes responsible for hydroxylation of JA reveals a missing step in JA metabolism, which is important for the inactivation of the hormone and subsequent down-regulation of JA-dependent defenses.

**SIGNIFICANCE STATEMENT:** In plants, the hormone jasmonic acid (JA) is synthesized in response to attack by pathogens and herbivores, leading to activation of defense responses. Rapidly following JA accumulation, the hormone is metabolized, presumably to prevent inhibitive effects of high JA levels on growth and development. The enzymes that directly inactivate JA were so far unknown. Here, we identify four jasmonate-induced oxygenases (JOXs) in Arabidopsis that hydroxylate jasmonic acid to form inactive 12-OH-JA. A mutant that no longer produces the four enzymes hyperaccumulates JA, exhibits reduced growth, and is highly resistant to attackers that are sensitive to JA-dependent defense. The JOX enzymes thus play an important role in determining the amplitude and duration of JA responses to balance the growth-defense tradeoff.

## INTRODUCTION

The lipid-derived phytohormone jasmonic acid (JA) is an essential signaling molecule in plant defense. In response to pathogen attack or wounding, JA levels accumulate resulting in activation of a subset of immune genes and the production of defensive secondary metabolites (1). Multiple negative feedback mechanisms control JA levels and JA-responsive gene expression, presumably to minimize inhibition of plant growth that is associated with JA-mediated defense responses (2). In healthy plants, under non-stressed conditions, low levels of JA are present, and the activation of JA-responsive genes is prevented by JAZ repressor proteins that bind transcriptional activators of the JA pathway (3–7). The conjugate of JA with isoleucine (JA-Ile) strongly promotes binding of JAZ repressors to the F-box protein COI1 (4, 8, 9), resulting in the degradation of JAZ proteins and subsequent activation of JA-responsive gene expression (10, 11). At the same time, *JAZ* gene expression is activated by JA resulting in the subsequent repression of JA-responsive gene expression (4, 5, 12). Excess JA and JA-Ile are also inactivated by hydroxylation, forming 12-hydroxy-JA (12-OH-JA) and 12-OH-JA-Ile (13–17).

The hydroxylated form of JA is thought to be inactive as it does not trigger degradation of JAZ repressors, and treatment with 12-OH-JA accordingly does not induce JA-responsive gene expression, nor does it inhibit root growth or seed germination (18, 19). 12-OH-JA has been identified in several plant species, including Arabidopsis, maize, potato, tomato and rice, and accumulates in Arabidopsis after wounding and *Botrytis cinerea* infection (13, 18, 20–22). Strikingly, hydroxylation of JA to 12-OH-JA by a monooxygenase produced by the blast fungus *Magnaporthe oryzae* attenuates plant immune responses to this pathogen (19). So far, no plant enzyme that hydroxylates JA has been identified (23). Hydroxylation of JA-Ile, on the other hand, has been described in Arabidopsis and is mediated by three cytochrome P450 enzymes, CYP94B3, CYP94B1 and CYP94C1 (Fig 1A, 14, 24–26). 12-OH-JA can be formed from 12-OH-JA-Ile by cleavage of the isoleucine group by two amidohydrolases (27, 28). However, an Arabidopsis double mutant that no longer produces these enzymes still accumulates 12-OH-JA, implying that other enzymes catalyze direct hydroxylation of JA to 12-OH-JA.

**Figure 1.**
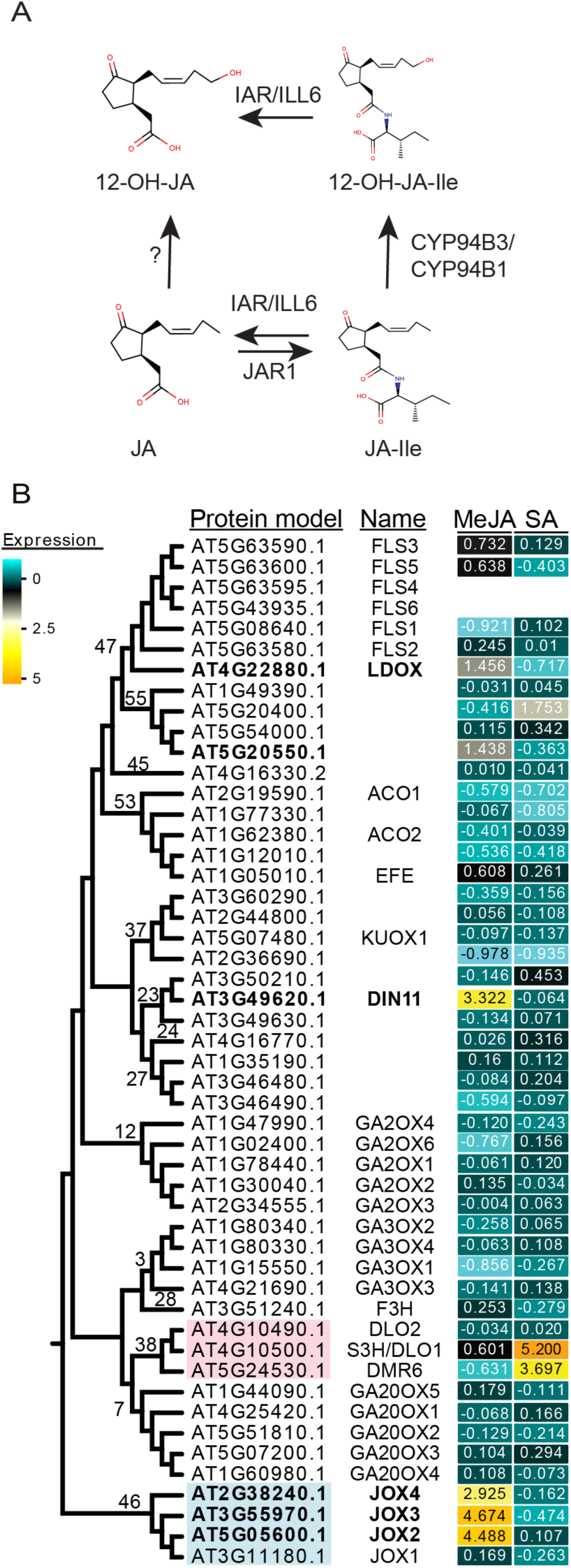
(A) Schematic of metabolism of JA and three JA-derived compounds: JA-Ile, 12-OH-JA, and 12-OH-JA-Ile. Enzymes that catalyze the conversions are indicated: JAR1 conjugates isoleucine to JA to form JA-Ile. CYP94B3 and CYP94C1 hydroxylate JA-Ile to 12-OH-JA-Ile. IAR and ILL6 can hydrolyze the Ile from JA-Ile or from 12-OH-JA-Ile, forming JA or 12-OH-JA, respectively. The enzyme hydroxylating JA to 12-OH-JA is hypothesized to be a 2OG oxygenase. **(B) Phylogenetic tree of SA- and MeJA-induced and related 2OG oxygenases of Arabidopsis.** The cladogram shows the relatedness of 50 2OG oxygenases selected from the phylogram in Figure S1. For each protein model, the name (when available) is supplied, and for each clade, the number assigned by Kawai et al. (2014) is indicated. The heat map indicates the log_2_ fold change of the corresponding genes in Arabidopsis seedlings 3 h after MeJA or SA treatment. Indicated in pink is clade 38, which contains SA-induced S3H/DLO1. 2OG oxygenases induced by MeJA are in bold. The JOX genes (clade 46) are indicated in blue.

Apart from cytochrome P450 enzymes, members of the 2-oxoglutarate (2OG) Fe(II)-dependent oxygenase family are involved in oxygenation/hydroxylation reactions in plants (29). Interestingly, several 2OG oxygenases were shown to hydroxylate and inactivate plant hormones; e.g. two different groups of 2OG oxygenases inactivate gibberellic acid (GA) by hydroxylating either bioactive 19-GA or an inactive precursor of GA (30, 31). More recently, the active form of auxin was reported to be hydroxylated and inactivated by the 2OG oxygenase DAO in rice (32) and Arabidopsis (33, 34). Finally, the defense hormone salicylic acid (SA) is hydroxylated by the 2OG oxygenase SA 3-HYDROXYLASE (S3H) (35). Since inactivation of hormones via hydroxylation by 2OG oxygenases is common in plants, we hypothesized that 2OG oxygenases could function as JA-hydroxylases as well. Here, we describe the identification of a clade of four 2OG oxygenases that are transcriptionally induced by JA, which we named JASMONATE-INDUCED OXYGENASES (JOX). We provide metabolic and biochemical evidence that these enzymes are responsible for hydroxylation of JA to 12-OH-JA.

Furthermore, phenotypic studies of mutant and overexpression lines show that the JOXs are involved in down-regulation of JA-dependent responses, thereby affecting plant defense and growth. These results identify a new class of enzymes in JA metabolism that perform an essential role in in controlling defense responses to necrotrophs and herbivorous insects.

## RESULTS

### Four JASMONATE-INDUCED OXYGENASES group in a distinct clade in Arabidopsis

We set out to investigate if JA-induced 2OG oxygenases could play a role in hydroxylation of JA to 12-OH-JA (Fig. 1A). First, a phylogenetic tree of 2OG oxygenases was constructed based on 93 Arabidopsis proteins that each contain two conserved 2OG oxygenase Pfam domains; the C-terminal PF03171 (2OG-Fe(II) oxygenase superfamily), and the N-terminal PF14226 (non-haem dioxygenase in morphine synthesis N-terminal). Phylogenetic clustering of the 93 proteins revealed distinct families (SI Appendix Fig. S1), that largely overlapped with previously described clades (29) which were based on six plant species ranging from unicellular alga to the flowering plants Arabidopsis and rice. Projection of transcriptome data showed that 2OG oxygenases that are induced in Arabidopsis seedlings by treatment with the defense-related hormones methyl jasmonate (MeJA) or SA were only present in a cluster of 50 proteins that encompass 14 clades as defined by Kawai *et al.* (29) (Fig. 1B). Clade 38 (in red) contains the *S3H/DLO1* and *DMR6* genes that are induced by SA (35, 36). The expression of six genes encoding 2OG oxygenases was induced more than 2-fold by MeJA treatment (indicated in bold; Fig. 1B). Of two weakly induced genes, At5g20550 is a gene of unknown function, whereas LDOX (At4g22880) encodes an enzyme involved in anthocyanin biosynthesis (37). A third MeJA-induced gene is *DIN11* (At3g49620), which was described as a senescence-associated gene responsive to viral infection (38). Strikingly, clade 46 (in blue) contains four 2OG-oxygenases of which 3 were clearly induced at 3 h after MeJA treatment (Fig. 1B).

The JA-responsiveness of the four 2OG oxygenase genes of clade 46 was experimentally verified in five-week-old plants treated with MeJA. This treatment increased transcript levels of all four genes (Fig. 2) that were, therefore, named *JASMONATE-INDUCED OXYGENASEs* (*JOXs*). The expression of *JOX1* (At3g11180), which was not MeJA-induced in seedlings according to publically available microarray data (Fig 1, 39), was highly induced in adults plants at 1 h after MeJA treatment, and was slightly higher than in mock-treated plants at 2-6 h after MeJA treatment (Fig. 2). The expression patterns of *JOX2* (At5g05600) and *JOX3* (At3g55970) were similar: induction was low at 1 h and 2 h after treatment, but high at 6 h after treatment. Expression of *JOX4* (At2g38240) was induced at all time points, but showed a different temporal behavior: it was highly induced at 1 h, lower at 2 h and high again at 6 h after treatment (Fig. 2). In the *coi1-1* mutant, that does not have a functional JA receptor, none of the *JOX* genes were induced by MeJA (Fig. 2). Interestingly, expression of the four *JOX* genes was strongly induced in plants infected by *B. cinerea* or infested by the caterpillar *Mamestra brassicae*, which both induce JA accumulation (SI Appendix Fig. S2). The fact that the four related *JOX* oxygenase genes are all activated by JA make them prime candidates to be involved in JA metabolism, similar to the SA-hydroxylase S3H/DLO1 that is transcriptionally induced by its substrate SA (Fig. 1B). As 2OG oxygenases are generally involved in hydroxylation and oxygenation reactions, we hypothesized that the JOX enzymes could hydroxylate JA.

**Figure 2.**
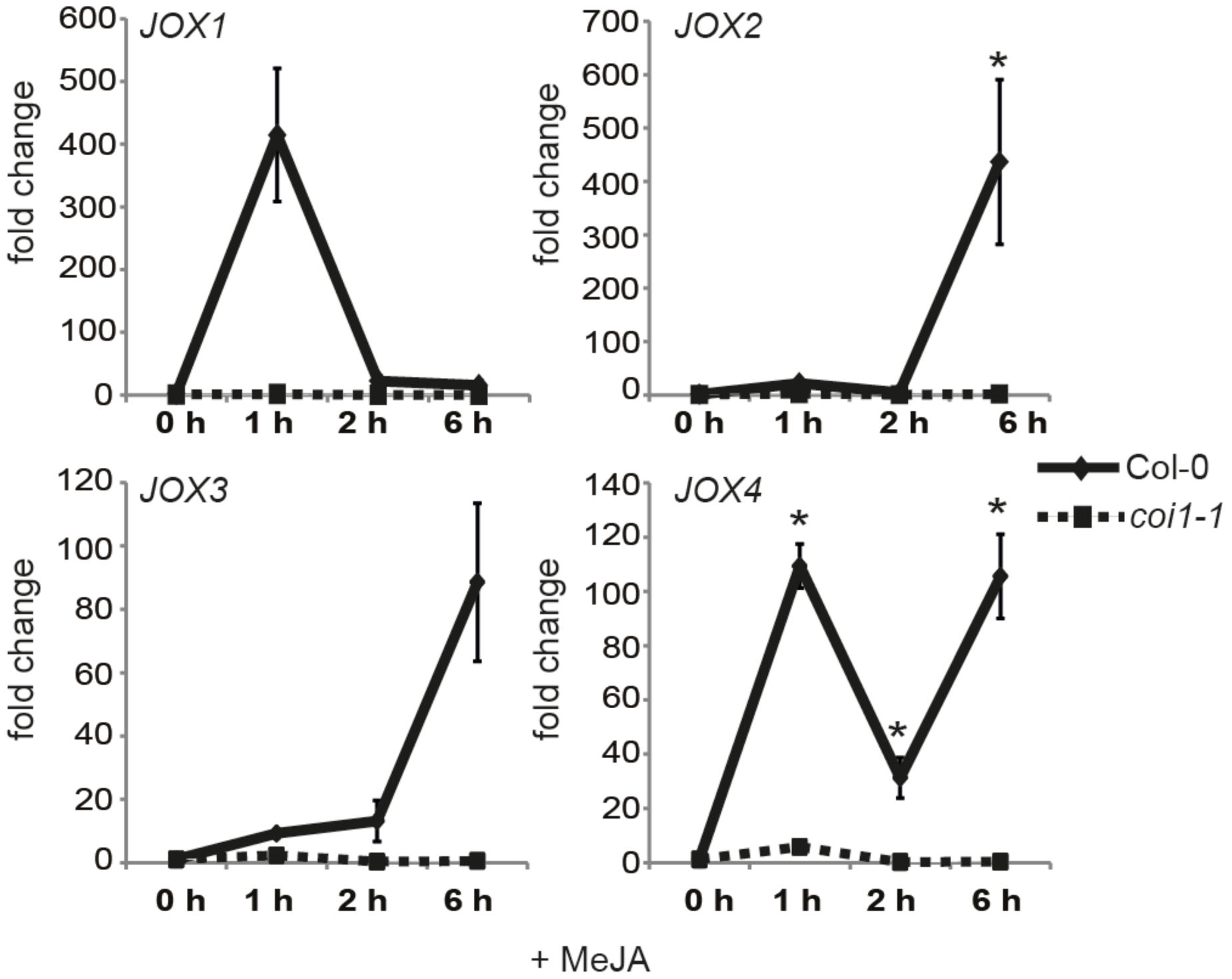
Expression of *JOX* genes after MeJA treatment. Expression analysis of *JOX1, JOX2, JOX3* and *JOX4* in response to MeJA in 5-week-old Col-0 and *coi1-1.* Shown is the expression (fold change) at 0 h, 1 h, 2 h, or 6 h after MeJA treatment, relative to that in mock-treated plants at the same time. Error bars indicate standard error. An asterisk indicates a significant higher expression compared to mock-treatment (*P* ≤ 0.05; two-way ANOVA).

**Figure 3.**
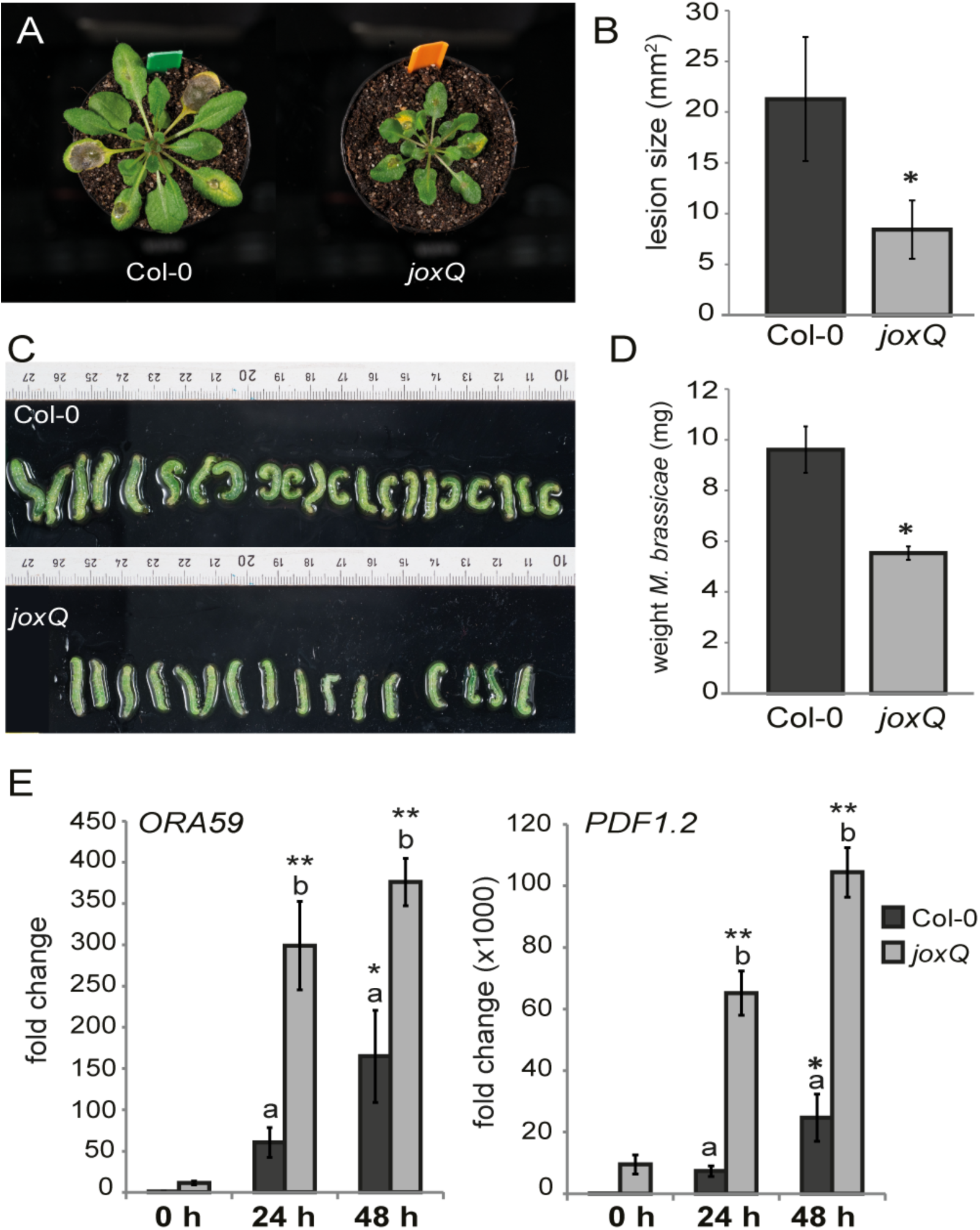
The *joxQ* quadruple mutant displays phenotypes reminiscent of activated JA signaling. **(A)** Representative pictures of reduced disease symptoms caused by *B. cinerea* infection on *joxQ* compared to Col-0. (**B)** Area size of necrotic lesions caused by *B. cinerea* in Col-0 and *joxQ*. **(C, D)** Size and weight of *M. brassicae* caterpillars on Col-0 and *joxQ* plants after 8 days of feeding. (B, D) An asterisk denotes a significant difference between Col-0 and *joxQ* (*t* test, *P*≤ 0.001). **(E)** Expression of the JA-responsive genes *ORA59* and *PDF1.2* before infection (0 h) and after 24 or 48 h of *B. cinerea* infection of Col-0 or *joxQ*, relative to Col-0 at 0 h. Different letters indicate significant differences between genotypes. An asterisk indicates a significant higher expression compared to Col-0 plants at 0 h (two-way ANOVA, Tukey post-hoc test; * *P*≤ 0.05; ** *P*≤ 0.001).

### JOX enzymes are **negative** regulators of JA responses

The presumed function of JOX enzymes in JA hydroxylation is expected to influence JA-related biological processes. We therefore set out to test phenotypes affected by JA in mutant plants. Since the four *JOX* genes could act redundantly, a quadruple mutant *jox1 jox2 jox3 jox4*, hereafter referred to as *joxQ*, was generated by crossing four T-DNA insertion lines in each of which one of the four *JOX* genes was disrupted. JA-related phenotypes, e.g. resistance to necrotrophic pathogens and herbivorous insects, were analyzed in the *joxQ* mutant. Lesions caused by infection with the necrotrophic fungus *B. cinerea* were significantly smaller in the *joxQ* mutant compared to those on wild-type Col-0 leaves (Fig. 3A&B). In addition, larvae of the generalist caterpillar *M. brassicae* were smaller and weighed significantly less when fed on *joxQ* compared to those fed on Col-0 (Fig. 3C&D). This suggests that defense against both necrotrophic pathogens and herbivorous caterpillars is upregulated in *joxQ* plants. To understand the molecular basis of the increased resistance in the *joxQ* mutant, we measured expression of JA-responsive defense genes before and after infection or infestation. Already before *B. cinerea* infection, expression of the JA/ET-responsive *PDF1.2* and *ORA59* genes was higher in the *joxQ* mutant compared to Col-0; the transcription factor gene *ORA59* was expressed 15-fold higher, while expression of *PDF1.2* was increased 9000-fold (Fig. 3E). During infection with *B. cinerea, ORA59* and *PDF1.2* levels strongly increased and were significantly higher in the *joxQ* mutant than in Col-0 (Fig. 3E). Similarly, expression of the JA-responsive transcription factor gene *MYC2* was higher in the *joxQ* mutant than in Col-0 under control conditions. Levels of this gene increased after *M. brassicae* feeding, and stayed slightly higher in the *joxQ* mutant (SI Appendix Fig. S3A).

JA is also known to inhibit plant growth and delay flowering time (40, 41). In accordance with this, the *joxQ* quadruple mutant was consistently smaller than wild-type plants (Fig. 3A) and exhibited reduced root growth on ½ MS medium (SI Appendix Fig. S3B). In addition, in the presence of 50 µM MeJA, the length of the main root was more strongly affected in *joxQ* (main root length 12.5% of untreated) than in Col-0 (20% of untreated) (SI Appendix Fig. S3B). These phenotypes support the idea that in wild-type plants JOX proteins suppress JA-mediated growth inhibition. Moreover, flowering time was delayed by 6 days in the *joxQ* mutant compared to Col-0 under short-day conditions and the mutant produced fewer seeds than wild-type Col-0 (SI Appendix Fig. S3C). The disease resistance and other JA-related phenotypes of the quadruple *joxQ* mutant are reminiscent of plants with activated JA responses, and are possibly caused by high JA levels.

### The *joxQ* mutant **hyperaccumulates** JA

The accumulation of JA could explain the phenotypes of the *joxQ* quadruple mutant and was therefore measured by LC-MS/MS (see SI Appendix Fig. S4 for standards). We found that JA levels were about 5x higher in the *joxQ* mutant than in Col-0 under non-treated conditions (Fig. 4A). Three hours after wounding, which is known to trigger JA accumulation, JA levels tripled to 18 ng/g FW in wild-type plants. In the *joxQ* mutant, JA rose to 146 ng/g FW (Fig. 4A). JA-Ile levels also increased after wounding in Col-0 and the *joxQ* mutant (Fig. 4A). The increased accumulation of JA in the *joxQ* mutant suggests that in wild-type plants the JOX proteins negatively affect the accumulation of JA, possibly via hydroxylation. To further study this, *JOX1* (At3g11180) was overexpressed in the Col-0 background (*JOX1* OX). In accordance with our hypothesis, JA and JA-Ile levels did not increase after wounding in this line (Fig. 4A). The results of both the mutant and overexpression line suggest that after wounding, the JOX enzymes act to reduce amounts of JA, possibly via hydroxylation.

**Figure 4.**
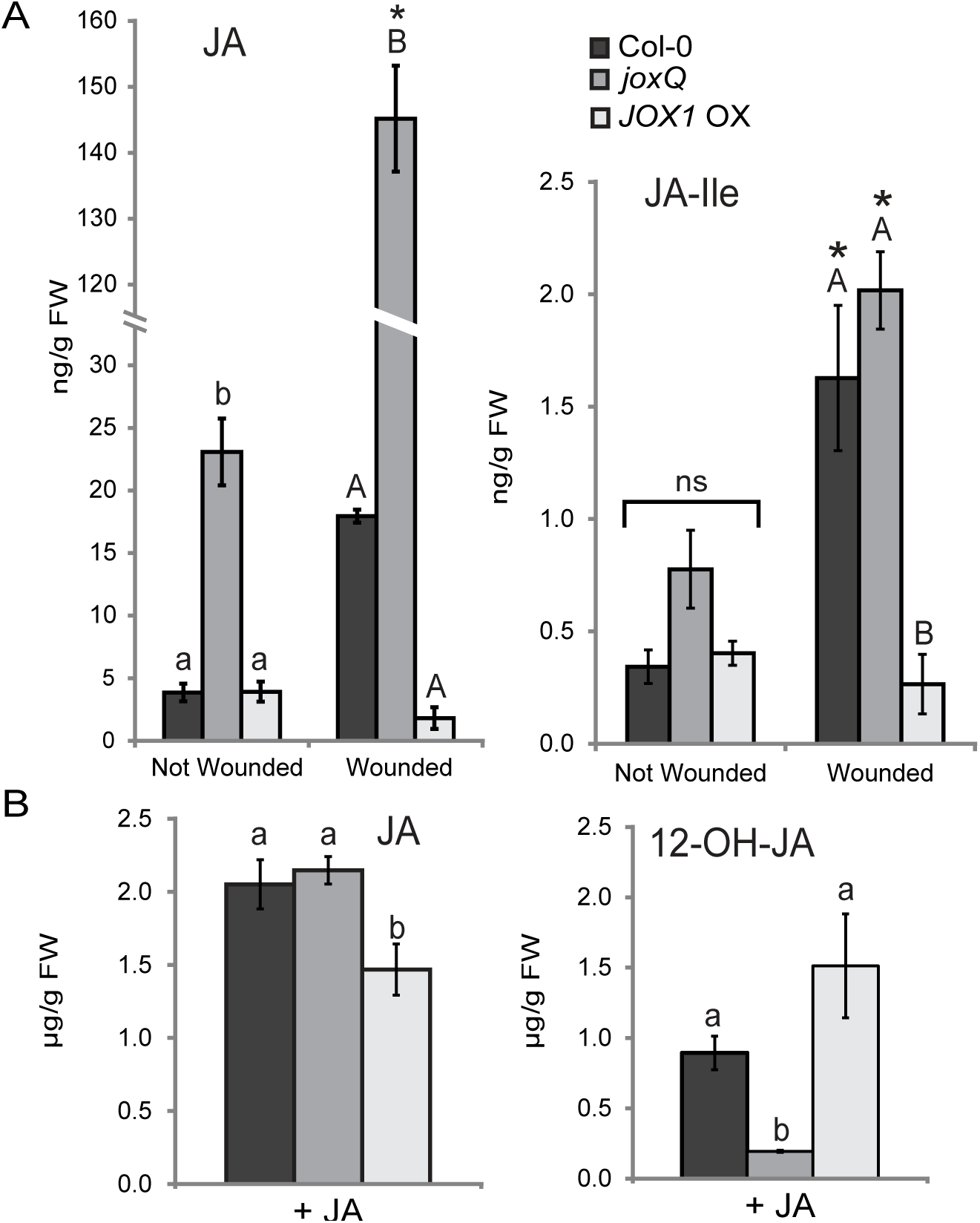
The *joxQ* mutant accumulates JA and is impaired in 12-OH-JA production after JA treatment. **(A)** JA and JA-Ile levels in leaves with or without wounding (3 h after mechanical damage) of Col-0, *joxQ* and *JOX1* OX plants. Each data point represents the mean of 4 biological replicates. Error bars indicate standard error. JA and JA-Ile levels were calculated by correcting for the internal standard and leaf weight. Different letters indicate statistically significant differences between genotypes at the same treatment, and an asterisk that wounding significantly induced the compound (two-way ANOVA; Tukey post-hoc test; *P*≤ 0.05). **(B)** Accumulation of JA and 12-OH-JA in plants exposed for 3 h to 100 µM JA. Each data point represents the mean of 4 biological replicates. Error bars indicate standard error. JA and 12-OH-JA levels were calculated by correcting for the internal standard and leaf weight. Different letters indicate statistically significant differences between genotypes (two-way ANOVA; Tukey post-hoc test; *P*≤ 0.05). All measured JA-related metabolites are shown in SI Appendix Table S2.

The hydroxylated form of JA, 12-OH-JA, was earlier shown to peak at 3 hours after wounding in wild-type Arabidopsis plants (20) and at this time point we measured 21 ng/g FW in Col-0 grown under our conditions (SI Appendix Fig. S5). If JOXs indeed hydroxylate JA, we would expect low levels of 12-OH-JA in the *joxQ* mutant. However, 12-OH-JA was about 4-fold higher in the *joxQ* mutant (84 ng/g FW) in response to wounding compared to Col-0 (SI Appendix Fig. S5 and Table S1). Nevertheless, the 12-OH-JA increase was lower than that of JA that was increased 8-fold in *joxQ* compared to Col-0. Possibly, 12-OH-JA detected in *joxQ* is generated by cleavage of the conjugated isoleucine of 12-OH-JA-Ile by aminohydrolases ILL6 and IAR3 (Fig 1A, 27). In *JOX1* OX we expected increased 12-OH-JA to accumulate, however, 12-OH-JA could not be detected in this line, even not after wounding (SI Appendix Fig. S5). We considered two hypotheses for this low 12-OH-JA level: (1) it is further metabolized and therefore not detectable, and/or, (2) it is not produced in sufficient amounts due to low levels of the JA substrate in the *JOX1* OX line. We did not find evidence for enhanced 12-OH-JA metabolism as we did not detect an increase in the downstream compounds 12-HSO_4_-JA, 12-OH-JA-Ile and 12-O-glycosyl-JA in *JOX1* OX plants (SI Appendix Table S1 and Fig. S5). In support of the low substrate levels, we find low JA (Fig. 4A), and reduced OPDA levels in five-week-old JOX1 OX plants (SI Appendix Table S1). As a positive feedback loop exists between JA levels and JA biosynthesis, we speculate that this feedback loop generates more JA in Col-0 than in JOX1 OX plants.

To reduce the effect of feedback mechanisms, we added equal amounts of exogenous JA to wild-type, *joxQ* and *JOX1* OX plants by immersing the leaves in a 100 µM JA solution and measured jasmonates three hours later. JA levels in treated leaves of *joxQ* were similar to those of Col-0, but lower in the *JOX1* OX line (Fig. 4B). Strikingly, the level of 12-OH-JA was much lower in the *joxQ* mutant than in Col-0. In contrast, in *JOX1* OX, the levels of 12-OH-JA were higher than in Col-0 (Fig. 4B). This indicates that the conversion of JA into 12-OH-JA is reduced in the *joxQ* mutant, and enhanced in the *JOX1* OX line. We conclude that the *joxQ* mutant hyperaccumulates JA and JA-Ile under basal and biotic stress conditions, and has reduced 12-OH-JA formation after treatment with exogenous JA. In contrast, a line overexpressing *JOX1* builds up less JA, and accumulates more of the hydroxylated form when treated with JA.

### Overexpression of each individual *JOX* complements *joxQ* phenotypes

The presumed metabolic function of JOX enzymes is to hydroxylate JA. To determine if all JOX enzymes have the capability to reduce JA levels and responses, we constitutively expressed each *JOX* gene from the 35S promoter in the quadruple *joxQ* mutant. For each *JOX*, at least two independent homozygous transformants were selected. First, we observed that overexpression of each *JOX* rescued the growth phenotype of the *joxQ* mutant. While five-week old *joxQ* plants were significantly smaller than Col-0, all *JOX*-transformed plants resembled Col-0 plants (Fig. 5A). Secondly, the high expression of *PDF1.2* that was observed in *joxQ* plants (Fig. 3E), was reverted to wild-type levels in all lines (Fig. 5B). Finally, we measured levels of JA and JA-derivatives in all transformed lines. In untreated adult plants of *joxQ*, JA levels were about 3 times higher than in Col-0. Overexpression of *JOX1*, *JOX2* and *JOX3* resulted in depletion of JA. Overexpression of *JOX4* resulted in JA levels similar to those in Col-0 (Fig. 5C). Taken together, these results show that each individual JOX can reduce JA levels to those in wild-type plants or lower, resulting in complementation of JA-related phenotypes (Fig. 5).

**Figure 5.**
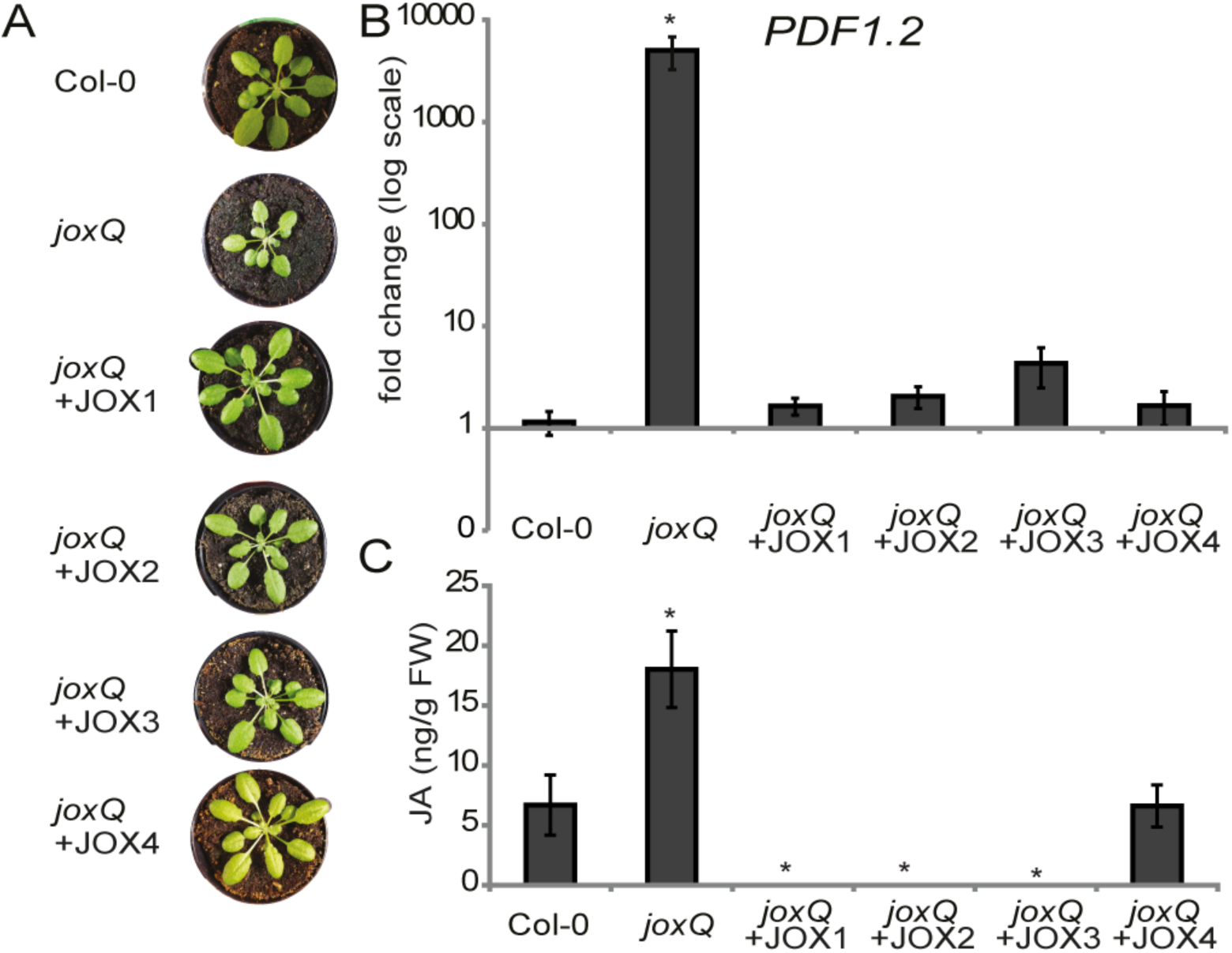
Overexpression of each individual JOX complements *joxQ* phenotypes. **(A)** Growth phenotype of representative plants of Col-0, *joxQ* and overexpression lines of JOX1, JOX2, JOX3, JOX4 in the *joxQ* background. **(B)** Expression of *PDF1.2* in untreated Col-0, *joxQ* and overexpression lines of JOX1, JOX2, JOX3, JOX4 in *joxQ*. Expression is normalized to the reference gene At1g13320, and relative to Col-0 plants. **(C)** Amount of JA in untreated leaves of Col-0, *joxQ* and overexpression lines of JOX1, JOX2, JOX3, JOX4 in *joxQ*. JA levels were calculated by correcting for the internal standard and leaf weight. (B, C) Each data point represents the mean of four to eight biological replicates. Error bars indicate standard error. For JOX overexpression lines, average is shown of two or three (JOX2) independent homozygous transformant lines. An asterisk denotes a significant difference between the genotype and Col-0 (*t* test, *P*≤ 0.05).

### JOX1, JOX2 and JOX4 hydroxylate JA *in vitro*

To determine whether the JOX proteins can catalyze the hydroxylation of JA to 12-OH-JA without other plant proteins present, assays were conducted with JA as substrate and recombinant JOX enzymes produced in *E. coli*. The reaction products were analyzed by LC-MS/MS. Lysates of *E. coli* expressing JOX1, JOX2 and JOX4 effectively converted JA to 12-OH-JA, resulting in almost complete conversion of the JA substrate, and high levels of 12-OH-JA (SI Appendix Fig. S6A). The lysate of *E. coli* expressing JOX3 produced low amounts of 12-OH-JA, which were higher than that of the negative control (reaction mixture without lysate) but not higher than the amount of 12-OH-JA produced in the enzymatic assay with *E. coli* expressing the SA-hydroxylase S3H/DLO1. We thus conclude that recombinant JOX1, JOX2 and JOX4 have JA-12-hydroxylase activity *in vitro* in the absence of other plant-derived enzymes and compounds.

## Discussion

Four JA-induced 2OG oxygenases (JOX) from a single paralogous family in Arabidopsis were identified that strongly contribute to negative regulation of JA-dependent responses. Plants in which the four genes were mutated (*joxQ*) accumulated high levels of JA, exhibited smaller growth, enhanced expression of defense genes, and resistance to both *B. cinerea* and *M. brassicae*. We provide evidence from metabolic measurements and enzymatic assays that JOX enzymes control JA levels by hydroxylation of JA to inactive 12-OH-JA. By keeping JA levels low the JOX enzymes are important in determining the amplitude and duration of JA responses and balance the growth-defense tradeoff.

Inactivation of plant hormones by 2OG oxygenase-mediated hydroxylation appears to be a common theme in plants to control their hormone levels. The hormones GA and SA are hydroxylated by 2OG oxygenases that are closely related to the JOXs, i.e. GA2OXs that hydroxylate GA (30) (clade 12 in Fig. 1B), and S3H (clade 38) that hydroxylates SA (35). Two more distantly related 2OG oxygenases were recently shown to inactivate the plant hormone auxin (33, 34). Clade 46 of the 2OG oxygenases as defined by Kawai *et al.* (29) consists of four Arabidopsis members, encoded by the genes At3g11180, At5g05600, At3g55970 and At2g38240, which we named *JOX1*, *JOX2*, *JOX3,* and *JOX4*, respectively. We show that the members of this clade act by oxidative inactivation of JA, and prevent overaccumulation of JA and indirectly its bio-active form JA-Ile.

The observation that JA-levels are high in the *joxQ* mutant already in untreated conditions (Fig. 4A), suggests that JOXs contribute to reduction of JA levels in unstressed growing conditions. Consequently, JA-responsive gene expression is higher in the *joxQ* plants than in Col-0 (Fig. 3E and S3A). JA-Ile is considered the biologically active form of JA as its binding to COI1–JAZ complexes leads to the degradation of JAZ repressor proteins and activation of JA-induced gene expression (4, 5, 9).However, we did not find increased JA-Ile levels in untreated *joxQ* plants. Possibly, JA-Ile is turned over so quickly that we could not detect it. It has been speculated that other derivatives of JA, or JA itself, can trigger gene activation. Interestingly, wound-induced expression of *MYC2 and PDF1.2* was not affected in the *jar1* mutant, which has reduced JA-Ile levels (12, 42). We also find increased expression of *MYC2* and *PDF1.2* in untreated *joxQ* plants. Further research should determine whether increased JA levels in the *joxQ* mutant cause these observed responses directly, or whether that occurs via JA-Ile or other JA-related metabolites.

After wounding, JA levels increased dramatically in the *joxQ* mutant. In wild type plants, JOX enzymes thus could function to reduce accumulation of JA. As JA biosynthesis genes are JA-responsive, it is likely that the higher JA levels in *joxQ* lead to increased biosynthesis and thus higher levels of JA. Supporting the idea that JA biosynthesis is upregulated in *joxQ*, we detected OPDA levels ∼4x times higher in the *joxQ* mutant than in Col-0. On the other hand, OPDA and JA levels are low in the *JOX1* overexpressing plants (Fig. 4A, SI Appendix Table S1). The reduced amount of JA substrate is possibly the cause that we do not find increased 12-OH-JA in *JOX1* OX plants. After adding exogenous JA to *JOX1* OX plants, more 12-OH-JA is generated compared to Col-0 (Fig. 4B), and in an enzymatic assay, JOX1 was able to form 12-OH-JA (SI Appendix Fig. S6). The JA hydroxylase activity of JOX1 is supported by these two experiments.

The induction of the *JOX* genes by MeJA treatment, by *M. brassicae* feeding and by *B. cinerea* infection, the first of which was shown to be dependent on JA-coreceptor COI1, shows that this JA-inactivating mechanism is controlled by the JA pathway itself. This is reminiscent of mechanisms of oxidative inactivation of hormones as discussed above and in particular of the mechanism of JA-Ile hydroxylation (14, 25). Enzymes from the cytochrome P450 family inactivate JA-Ile: CYP94B1 and CYP94B3 hydroxylate JA-Ile to 12-OH-JA-Ile (24–26) and CYP94C1 further oxygenates this compound to 12-COOH-JA-Ile (14). Our identification of four 2OG oxygenases that convert JA into the inactive 12-OH-JA, further elucidates JA metabolism. From 12-OH-JA-Ile, 12-OH-JA can be produced by the amidohydrolases IAR3 and ILL6, which cleave the isoleucine group of JA-Ile and 12-OH-JA-Ile (Fig 1A, 27). However, the JOX enzymes can hydroxylate JA directly, and majorly contribute to the removal of JA in plants in undisturbed growth conditions and in response to wounding or pathogen attack. The quadruple mutant of these enzymes shows similar phenotypes as the JA-Ile hydroxylase mutants, e.g. enhanced expression of JA-responsive genes and increased sensitivity to JA-dependent inhibition of root growth (Fig 3, 14). This suggests that hydroxylation of JA by the JOX enzymes contributes to inactivation of the active JA signal to a similar extent as hydroxylation of JA-Ile. The levels of bioactive JA-Ile are likely directly influenced by the amount of JA, as we show that levels of JA and JA-Ile are in equilibrium after wounding (Fig. 4A).

The importance of hydroxylation of JA is emphasized by the apparent evolution of four different enzymes with the same function. In an enzyme assay, recombinant JOX1, JOX2 and JOX4 were shown to be able to produce 12-OH-JA from JA in the absence of other plant proteins. JOX3 was unable to hydroxylate JA in this *in vitro* assay. However, in our complementation studies, all four enzymes, including JOX3, were shown to be able to complement all phenotypes of the *joxQ* mutant, indicating that each JOX enzyme can lower JA levels *in vivo.* Possibly, JOX3 requires activation by another plant protein or compound before it is active as an JA-hydroxylase, or folding of the recombinant JOX3 protein is not correct. JOX proteins are found in a broad taxonomic range of multicellular plant species (29) but the number of JOX orthologs differs per species. Interestingly, the monocot and dicot JOX orthologs group in separate phylogenetic branches, suggesting that ancestral flowering plants had a single *JOX* gene, while extant species have 2-4 paralogs. It is possible that each paralogous enzyme functions in a different process in the plant or in distinct plant tissues. Preliminary evidence for this comes from the timing and amplitude of the expression induced by MeJA, *B. cinerea* and *M. brassicae,* which was different between the *JOX* genes (Fig. 2, SI Appendix Fig. S2). So far, we have only tested expression in leaf tissue, while spatial expression of each *JOX* gene within the leaf or within the plant could be different. Similarly, the SA-hydroxylase gene *DLO1* and its paralog *DMR6* have similar but distinct activities due to their pathogen-induced expression in different parts of downy mildew-infected leaves (36). Experiments to localize the expression of *JOXs* and complementation assays under their own promoter could further elucidate the different functionalities of the 4 JOX genes.

In plants, JA is converted to derivatives that are biologically active, reduced active, or inactive (16). 12-OH-JA has been characterized as an inactive form of JA, as it is not capable of degrading JAZ9 and does not induce expression of JA-responsive genes or inhibition of root growth (15, 18, 19). The compound has been suggested to have tuber-inducing capabilities (22), but what the role of this compound is in non-tuber forming plants such as Arabidopsis is not clear. It is likely that hydroxylation of JA is a quick mechanism to inactivate JA. Following production of JA, expression of *JOX* genes would quickly be induced, after which the accumulation of JA, and subsequently the expression JA-responsive genes, is dampened. This is yet another negative feedback system that would be active in the JA pathway, in addition to, for example, the activation of JAZ repressor genes by JA (4, 5, 12). Moreover, similar hydroxylation mechanisms control levels of other plant hormones (25, 33, 35). The identification of JOX enzymes elucidates another major step in the metabolic pathway of plants. Our data shows that it is imperative that plants balance JA levels by controlling JA metabolism, as inhibitive effects on growth are evident in the *joxQ* mutant. As expression of the JOX genes is induced by JA, plants can quickly shut off JA-dependent responses after their activation to prevent negative effect of high JA levels on growth and development. The JOX enzymes thus contribute to balance the growth/defense tradeoff.

## Material and Methods

Arabidopsis genes encoding 2OG-oxygenases were selected from Biomart (plant.ensembl.org/biomart), aligned, processed for phylogenetic analysis, and plotted with gene expression data as described in SI Appendix Material and Methods. The generation of the quadruple *jox* mutant and *JOX* OX lines, as well as their phenotypic characterization and chemical profiles is detailed in SI Appendix Material and Methods, as well as the enzymatic assays on recombinant JOX enzymes produced in *E. coli*.

## Acknowledgments

We thank Léon Westerd for rearing of *M. brassicae*, Hans van Pelt for taking photographs of plants and caterpillars, and Tom Raaymakers for help with the figures. This work was partly financed by the Netherlands Organization for Scientific Research (NWO) through the Dutch Technology Foundation (STW) STW VIDI Grant no. 11281 (to SCMvW). We are grateful to Enza Zaden B.V. for supporting the initial research on the *JOX* genes.

## References

1. Campos ML, Kang JH, Howe GA (2014) Jasmonate-triggered plant immunity. J Chem Ecol 40(7):657–675.

2. Huot B, Yao J, Montgomery BL, He SY (2014) Growth-defense tradeoffs in plants: a balancing act to optimize fitness. Mol Plant 7(8):1267–1287.

3. Zhu Z, et al. (2011) Derepression of ethylene-stabilized transcription factors (EIN3/EIL1) mediates jasmonate and ethylene signaling synergy in Arabidopsis. Proc Natl Acad Sci U S A 108(30):12539–12544.

4. Thines B, et al. (2007) JAZ repressor proteins are targets of the SCFCOI1 complex during jasmonate signalling. Nature 448(7154):661–665.

5. Chini A, et al. (2007) The JAZ family of repressors is the missing link in jasmonate signalling. Nature 448(7154):666–671.

6. Fernández-Calvo P, et al. (2011) The Arabidopsis bHLH transcription factors MYC3 and MYC4 are targets of JAZ repressors and act additively with MYC2 in the activation of jasmonate responses. Plant Cell 23(2):701–715.

7. Niu Y, Figueroa P, Browse J (2011) Characterization of JAZ-interacting bHLH transcription factors that regulate jasmonate responses in Arabidopsis. J Exp Bot 62(6):2143–2154.

8. Sheard LB, et al. (2010) Jasmonate perception by inositol-phosphate-potentiated COI1-JAZ co-receptor. Nature 468(7322):400–405.

9. Fonseca S, et al. (2009) (+)-7-iso-Jasmonoyl-L-isoleucine is the endogenous bioactive jasmonate. Nat Chem Biol 5(5):344–350.

10. Devoto A, et al. (2002) COI1 links jasmonate signalling and fertility to the SCF ubiquitin-ligase complex in Arabidopsis. Plant J 32(4):457–466.

11. Katsir L, Schilmiller AL, Staswick PE, He SY, Howe GA (2008) COI1 is a critical component of a receptor for jasmonate and the bacterial virulence factor coronatine. Proc Natl Acad Sci U S A 105(19):7100–7105.

12. Chung HS, et al. (2008) Regulation and function of Arabidopsis JASMONATE ZIM-domain genes in response to wounding and herbivory. Plant Physiol 146(3):952–964.

13. Seto Y, et al. (2009) Purification and cDNA cloning of a wound inducible glucosyltransferase active toward 12-hydroxy jasmonic acid. Phytochemistry 70(3):370–379.

14. Heitz T, et al. (2012) Cytochromes P450 CYP94C1 and CYP94B3 catalyze two successive oxidation steps of plant hormone jasmonoyl-isoleucine for catabolic turnover. J Biol Chem 287(9):6296–6306.

15. Gidda SK, et al. (2003) Biochemical and molecular characterization of a hydroxyjasmonate sulfotransferase from Arabidopsis thaliana. J Biol Chem 278(20):17895–17900.

16. Wasternack C, Strnad M (2016) Jasmonate signaling in plant stress responses and development - active and inactive compounds. N Biotechnol 33(5):604–613.

17. Guranowski A, Miersch O, Staswick PE, Suza W, Wasternack C (2007) Substrate specificity and products of side-reactions catalyzed by jasmonate:amino acid synthetase (JAR1). FEBS Lett 581(5):815–820.

18. Miersch O, Neumerkel J, Dippe M, Stenzel I, Wasternack C (2008) Hydroxylated jasmonates are commonly occurring metabolites of jasmonic acid and contribute to a partial switch-off in jasmonate signaling. New Phytol 177(1):114–127.

19. Patkar RN, et al. (2015) A fungal monooxygenase-derived jasmonate attenuates host innate immunity. Nat Chem Biol 11(9):733–740.

20. Glauser G, et al. (2008) Spatial and temporal dynamics of jasmonate synthesis and accumulation in Arabidopsis in response to wounding. J Biol Chem 283(24):16400–16407.

21. Aubert Y, Widemann E, Miesch L, Pinot F, Heitz T (2015) CYP94-mediated jasmonoyl-isoleucine hormone oxidation shapes jasmonate profiles and attenuates defence responses to Botrytis cinerea infection. J Exp Bot 66(13):3879–3892.

22. Yoshihara T, et al. (1989) Structure of a Tuber-inducing Stimulus from Potato Leaves (Solanum tuberosum L.). Agric Biol Chem 53(10):2835–2837.

23. Koo AJ, Howe GA (2012) Catabolism and deactivation of the lipid-derived hormone jasmonoyl-isoleucine. Front Plant Sci 3:19.

24. Koo AJ, et al. (2014) Endoplasmic reticulum-associated inactivation of the hormone jasmonoyl-L-isoleucine by multiple members of the cytochrome P450 94 family in Arabidopsis. J Biol Chem 289(43):29728–29738.

25. Koo AJ, Cooke TF, Howe GA (2011) Cytochrome P450 CYP94B3 mediates catabolism and inactivation of the plant hormone jasmonoyl-L-isoleucine. Proc Natl Acad Sci U S A 108(22):9298–9303.

26. Kitaoka N, et al. (2011) Arabidopsis CYP94B3 encodes jasmonyl-L-isoleucine 12-hydroxylase, a key enzyme in the oxidative catabolism of jasmonate. Plant Cell Physiol 52(10):1757–1765.

27. Widemann E, et al. (2013) The amidohydrolases IAR3 and ILL6 contribute to jasmonoy-l isoleucine hormone turnover and generate 12-hydroxyjasmonic acid upon wounding in Arabidopsis leaves. J Biol Chem 288(44):31701–31714.

28. Bhosale R, et al. (2013) Predicting gene function from uncontrolled expression variation among individual wild-type Arabidopsis plants. Plant Cell 25(8):2865–2877.

29. Kawai Y, Ono E, Mizutani M (2014) Evolution and diversity of the 2-oxoglutarate-dependent dioxygenase superfamily in plants. Plant J 78(2):328–343.

30. Rieu I, et al. (2008) Genetic analysis reveals that C19-GA 2-oxidation is a major gibberellin inactivation pathway in Arabidopsis. Plant Cell 20(9):2420–2436.

31. Schomburg FM, Bizzell CM, Lee DJ, Zeevaart JA, Amasino RM (2003) Overexpression of a novel class of gibberellin 2-oxidases decreases gibberellin levels and creates dwarf plants. Plant Cell 15(1):151–163.

32. Zhao Z, et al. (2013) A role for a dioxygenase in auxin metabolism and reproductive development in rice. Dev Cell 27(1):113–122.

33. Zhang J, et al. (2016) DAO1 catalyzes temporal and tissue-specific oxidative inactivation of auxin in Arabidopsis thaliana. Proc Natl Acad Sci U S A 113(39):11010–11015.

34. Porco S, et al. (2016) Dioxygenase-encoding AtDAO1 gene controls IAA oxidation and homeostasis in Arabidopsis. Proc Natl Acad Sci U S A 113(39):11016–11021.

35. Zhang K, Halitschke R, Yin C, Liu CJ, Gan SS (2013) Salicylic acid 3-hydroxylase regulates Arabidopsis leaf longevity by mediating salicylic acid catabolism. Proc Natl Acad Sci U S A 110(36):14807–14812.

36. Zeilmaker T, et al. (2015) DOWNY MILDEW RESISTANT 6 and DMR6-LIKE OXYGENASE 1 are partially redundant but distinct suppressors of immunity in Arabidopsis. Plant J 81(2):210–222.

37. Shan X, Zhang Y, Peng W, Wang Z, Xie D (2009) Molecular mechanism for jasmonate-induction of anthocyanin accumulation in Arabidopsis. J Exp Bot 60(13):3849–3860.

38. Fernández-Calvino L, et al. (2016) Activation of senescence-associated Dark-inducible (DIN) genes during infection contributes to enhanced susceptibility to plant viruses. Mol Plant Pathol 17(1):3–15.

39. Goda H, et al. (2008) The AtGenExpress hormone and chemical treatment data set: experimental design, data evaluation, model data analysis and data access. Plant J 55(3):526–542.

40. Zhai Q, et al. (2015) Transcriptional Mechanism of Jasmonate Receptor COI1-Mediated Delay of Flowering Time in Arabidopsis. Plant Cell 27(10):2814–2828.

41. Zhang Y, Turner JG (2008) Wound-induced endogenous jasmonates stunt plant growth by inhibiting mitosis. PLoS ONE 3(11):e3699.

42. Suza WP, Staswick PE (2008) The role of JAR1 in Jasmonoyl-L: -isoleucine production during Arabidopsis wound response. Planta 227(6):1221–1232.

